# Nitrogen Availability and Utilisation of Oligopeptides by Yeast in Industrial Scotch Grain Whisky Fermentation

**DOI:** 10.1101/2024.07.02.601656

**Authors:** Hidde Yaël Berg, Georg Arju, Ildar Nisamedtinov

## Abstract

Scotch grain whisky is produced with a substantial proportion of unmalted grains, which can result in nitrogen deficiency for yeast in the fermentable grain mash. This study examined nitrogen source availability and utilisation by three commercial whisky strains (*Saccharomyces cerevisiae*) during Scotch grain whisky fermentation, focusing on oligopeptides. Peptide uptake kinetics in synthetic whisky mash with defined peptides showed that oligopeptides of up to nine amino acids were taken up by the strains, albeit with some variability between the strains. The study found that peptides with appropriate molecular weights could replace free amino acids without negatively affecting fermentation kinetics. Moreover, fermentation performance improved when additional nitrogen was provided via peptides rather than diammonium phosphate. Analysis of industrial grain mash indicated that despite low initial yeast assimilable nitrogen, residual proteolytic activity from malt increased nitrogen availability during fermentation. Approximately 30% of the nitrogen consumed by yeast during grain mash fermentation was derived from peptides. LC-HRMS peptide analysis revealed complex dynamics of peptide formation, degradation, and utilisation. This study highlights the importance of oligopeptides in ensuring optimal fermentation efficiency in Scotch grain mash and similar substrates.

## Introduction

A readily available supply of easily assimilable nitrogen compounds, including amino acids and low-molecular-weight oligopeptides, is required for *Saccharomyces cerevisiae* to attain optimal fermentation efficiency during whisky fermentation ^[1–3]^. Assimilated amino acids and peptides can play both anabolic and catabolic roles in yeast, for example, through the biosynthesis of structural and functional proteins and nucleic acids ^[4, 5]^ or the formation of higher alcohols and their associated esters that impart flavour properties of the final product ^[6, 7]^. A prevalent belief is that most of the assimilable nitrogen for yeast in wort is produced during the malting process ^[8, 9]^. Unlike the production process of malt whisky, which the whole mash bill is composed of malted grains, Scotch grain whisky worts are primarily derived from unmalted grains ^[10]^. Consequently, it is crucial to consider a potential limitation of nitrogenous compounds in this process.

Scotch grain whisky is usually produced from mash containing approximately 90% (w/w) unmalted grains, most commonly wheat ^[8, 10]^. The unmalted grains are mechanically milled to release starch from the grain’s protein matrix and then mixed with process liquor, which is a combination of process water, backset, distillation condensate, and weak worts ^[9]^. The resulting grain slurry is subsequently cooked to gelatinise the starch. The cooking temperature varies based on the specific gelatinisation temperature of the starch. Nevertheless, most Scotch grain distilleries typically use much higher temperatures (95-145°C) for starch gelatinisation ^[9]^. The cooked grain slurry is then combined with milled malt slurry in a conversion tank and maintained at mashing temperatures (63-65°C) for up to 30 min ^[9]^. The primary objective of this process, termed conversion, is to transform gelatinised starch into fermentable sugars (mostly maltose, glucose, and maltotriose) ^[13]^. Traditionally, wort has been separated from the grains to provide a clear fermentable liquid. However, the wort separation step has been largely eliminated in modern grain whisky distilleries; instead, the entire mash is transferred to the fermenter after cooling and pitching with yeast ^[9]^.

As mentioned above, it is widely considered that a significant proportion of the yeast-assimilable nitrogen present in the final wort is derived from the malt. During the malting process, proteolytic enzymes are released into the endosperm to degrade grain storage proteins (i.e. hordeins, glutelins, albumins, and globulins) into peptides and free amino acids ^[11]^. Many of these enzymes, including aminopeptidases and some endoproteinases, are sensitive to heat and are largely inactivated at temperatures above 55°C, used during the conversion step ^[12, 13]^. However, studies have described the formation of soluble nitrogenous compounds during conversion by heat-stable endoproteinases and carboxypeptidases ^[13, 14]^. These proteases are inactivated only at temperatures above 70°C ^[13]^. Unlike brewing, the malt stream in the Scotch grain whisky production process does not reach this temperature, nor does it contain a separation and boiling step ^[9, 10, 15]^. Consequently, in addition to the conversion step, the proteolytic activity can also be present during fermentation. In the context of Scotch grain whisky production, where the malted substrate constitutes on average 10% of the mash bill, this may be crucial for supplying sufficient nitrogen for yeast to ensure good fermentation kinetics, especially as the addition of exogenous proteolytic enzymes or nitrogen supplements is prohibited ^[16]^.

The utilisation of oligopeptides as a yeast nitrogen source has received limited attention in the field of Scotch grain whisky production. Our previous study on wine yeast showed that oligopeptides can supply yeast with sufficient nitrogen to complete fermentation without the need for additional amino acids ^[16]^. Thus, oligopeptides may also be an important secondary source of nitrogen for yeast during Scotch grain whisky fermentation.

In this study, we aim to investigate the availability, production, and utilisation of different yeast-assimilable nitrogen sources during Scotch grain whisky fermentation. Three commercial strains of *Saccharomyces cerevisiae*, commonly used in whisky production, were selected for analysis. The focus was on the role of oligopeptides as a secondary source of nitrogen for yeast. Our findings provide insights into nitrogen availability in Scotch grain whisky fermentation and highlight the importance of oligopeptides for ensuring optimal fermentation efficiency.

## Materials and Methods

### Strains

Three commercial *S. cerevisiae* whisky yeast strains (denoted as SC1, SC2 and SC3) were provided by Lallemand Inc. (Montreal, QC, Canada). The strains were stored as cryocultures in 25% glycerol at -80 ^°^C. The relevant strain information is presented in Table 1.

**Table 1.** Whisky strains used in this study. The information displayed in the table has been summarised from the provided technical data sheets of the respective strains (Lallemand Inc., Montreal, QC, Canada).

| Strain acronym | Type of whisky recommended for | Relative nitrogen requirement* |
| --- | --- | --- |
| SC1 | Grain Whisky | Low |
| SC2 | Grain and malt whisky | Low |
| SC3 | Malted grain whisky | High |

\*Nitrogen requirement relative to the other strains used in this study

Each yeast inoculum was prepared by streaking the cryoculture on a yeast peptone dextrose (YPD) agar plate (1% yeast extract, 2% bacteriological peptone, and 2% glucose) and pitching a single colony into a shaker flask containing YPD broth. The flasks were incubated overnight at 30°C, with shaking at 150 rpm. Cells were washed twice with equal volumes of sterile 0.9% NaCl prior to inoculation into fermenters to deliver 5 × 10^6^ cells/mL, based on OD600 measurements.

### Fermentations

#### Synthetic whisky mash fermentations

Fermentation experiments were performed on synthetic whisky mash (SWM) to confirm the importance of peptides as a nitrogen source to whisky yeast. In addition, the potential benefits of nitrogen supplementation, either through organic or inorganic means, were investigated.

#### Preparation of the bovine serum albumin hydrolysate

A bovine serum albumin (BSA) enzymatic hydrolysate was used as a source of peptides in a synthetic whisky mash (SWM). The hydrolysate was prepared according to a previously reported method ^[16, 17]^.

The hydrolysate was produced from BSA (40 g/L) in a 1 L benchtop fermenter (Applikon, Delft, The Netherlands) using the industrial protease COROLASE^®^7089 (AB Enzymes, Darmstadt, Germany). The protease dose rate was 0.5% w/w BSA, and hydrolysis was performed at pH 7 (maintained by titration using 2 M NaOH) for 20 h at 50 °C. After completion of BSA hydrolysis, the hydrolysate was filtered with a 10 kDa Vivaflow^®^ 200 10,000 MWCO Hydrosart crossflow cassette (Sartorius, Göttingen, Germany) to remove the protease and larger peptides. To further concentrate the smaller MW peptide fraction (≤ 2 kDa), another filtration step was performed using a Vivaflow^®^ 200 2,000 MWCO Hydrosart crossflow cassette (Sartorius, Göttingen, Germany). The hydrolysate was then freeze-dried and stored at –20 °C until further use.

#### Synthetic whisky mash (SWM) composition

The composition of SWM is an adaptation of the synthetic grape must described by Salmon and Barre ^[18]^. Citric and malic acids were excluded from the medium, and the sugar concentration, free amino acid ratios (Table S1), and pH were adjusted to mimic whisky mash. These values were based on measurements taken from a commercial whisky production process.

The base SWM contained 150 g/L maltose and glucose (9:1 w/w) and was modified by adding different nitrogen sources (Table 2). Thus, in the first experiment (AP230), 130 mg/L of nitrogen was added with amino acids, and 100 mg/L was added in the form of small molecular weight peptides (smaller than 2 kDa) from the BSA hydrolysate. In the second experiment, all nitrogen (200 mg/L) was supplied in the form of small-MW peptides from the BSA hydrolysate (P230). To study the effect of additional nitrogen supplementation with (in)organic sources, 70 mg/L of nitrogen was added to the P230 medium from either the BSA hydrolysate (PP300) or diammonium phosphate (PD300). The pH of all the media was adjusted to 5.2. The media were then filter-sterilised using a 0.22 μm Steritop^®^ vacuum-driven disposable filtration system (Merck-Millipore, Burlington, MA, USA) prior to yeast inoculation.

**Table 2.**
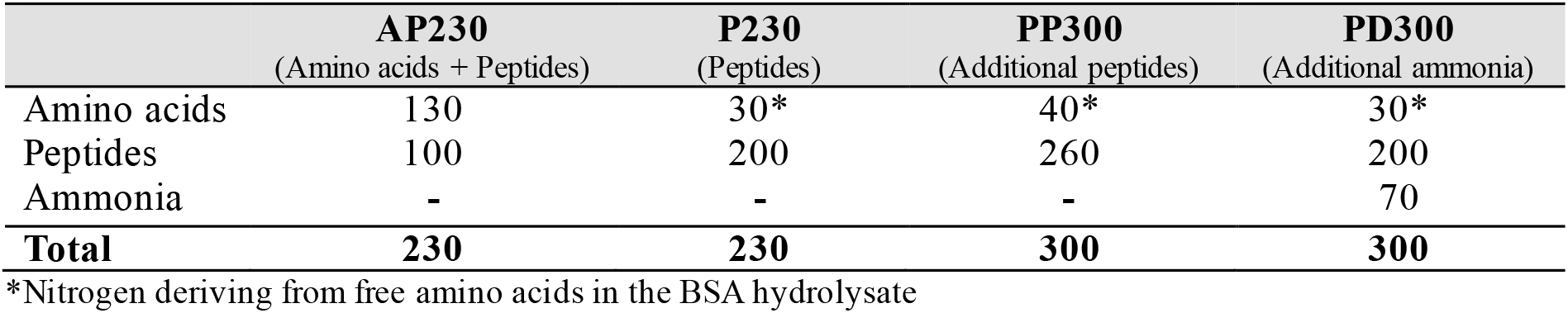
Nitrogen composition of the synthetic media used in this study. All media contained 150 g/Lmaltose and glucose (9:1, w/w). All values are displayed as milligrams of nitrogen per litre. Free proline was not taken into account as it is not or poorly assimilated by yeasts during fermentation ^[4, 40]^.

|  | AP230<br>(Amino acids + Peptides) | P230<br>(Peptides) | PP300<br>(Additional peptides) | PD300<br>(Additional ammonia) |
| --- | --- | --- | --- | --- |
| Amino acids | 130 | 30* | 40* | 30* |
| Peptides | 100 | 200 | 260 | 200 |
| Ammonia | - | - | - | 70 |
| **Total** | **230** | **230** | **300** | **300** |

\*Nitrogen deriving from free amino acids in the BSA hydrolysate

#### Fermentations

All fermentations on synthetic media (100 mL) were performed in triplicate at 24 °C in 100 mL Pyrex™ bottles equipped with a GL45 open-top PBT screw cap and PYREX™ Media Bottle Septum (Corning Inc., Corning, NY, USA). A gas outlet was installed to prevent overpressure by piercing the septum with a Sterican^®^ Ø 0.8 × 40 mm single-use hypodermic needle (B. Braun, Melsungen, Germany) attached to a Millex-FG 0.2 µm hydrophobic PTFE filter (Merck KGaA, Darmstadt, Germany).

Samples (2 mL) were collected every 6 h for the first 48 h, every 24 h until 120 h, and every 48 h until 192 h. Monitoring of CO_2_ production was continued until the endpoint (288 h), after which a final sample was collected. The biomass density of the samples was assessed by measuring the optical density at 600 nm using an Ultrospec^®^ 10 Cell Density Meter (Biochrom, Ltd., Cambridge, UK). The specific production rate of CO_2_, monitored gravimetrically, was used as the main indicator of the fermentation progress. After centrifugation at 9600 × g for 10 min at 4 °C in a MicroCL 21R Microcentrifuge (Thermo Fisher Scientific, MA, USA), the supernatant was filtered through Minisart RC 0.2 μm syringe filters (Sartorius, Göttingen, Germany) and stored at -20 °C until further analysis.

### Grain mash fermentations

Unfiltered grain mash, produced from approximately 10% malted barley and 90% unmalted wheat, was obtained from a commercial Scotch grain whisky distillery. Fermentation was performed in 500 mL of grain mash in 1 L Biobundle benchtop fermenters (Applikon, Delft, the Netherlands) using the cultivation control software BIOEXPERTV2 v2.96 (Applikon, Delft, the Netherlands). The fermenters were equipped with pH and temperature sensors. The production of CO_2_ was measured with an HPR-20 R&D Benchtop Gas Analysis System (Hiden Analytical Ltd., Warrington, UK) using MASsoft 10 Professional software (Hiden Analytical Ltd., Warrington, UK). To allow for stable measurement of CO_2_ production, the headspace was flushed at a rate of 100 ml/min (5% CO _2_, 95% N_2_).

The fermentation temperature was raised from 25 °C to 34 °C in the first 24 h to mimic the heat production that occurs during the factory process due to yeast activity, and then maintained at 34 °C for 60 h. All fermentations were performed in triplicate. Samples (10 mL) were collected every 12 h from the bottom of the fermenter using a sample port and a 10 ml sterile syringe (B. Braun, Melsungen, Germany).

The formation of free amino acids and peptides due to residual proteolytic activity in the grain mash was analysed in an uninoculated grain mash by simulating the changes in temperature, pH, and ethanol concentration during fermentation. The change gradients of the latter parameters were based on the actual measurements of grain mash fermentation with strain SC2 and are shown in

Table **3**. The uninoculated mash contained 200 μg/mL G418 (InvivoGen, Toulouse, France) and 2 ppm Lactoside 247™ (Lallemand, Montreal, QC, Canada) to prevent microbial growth.

**Table 3.** Temperature, pH, and ethanol concentration gradients used in uninoculated grain mash to study free amino acid and peptide production by native malt proteases. The ethanol concentration was increased stepwise by injecting 96% ethanol into the bioreactor.

| Time (h) | Temperature (°C)<br>(Gradient) | Ethanol (g/L)<br>(Manual every 12h) | pH<br>(Gradient) |
| --- | --- | --- | --- |
| 0 | 25 | 0 | 5.2 |
| 12 | 29.5 | 20 | 4.75 |
| 24 | 34 | 50 | 4.3 |
| 36-60 | 34 | 70 | 4.3 |

### Analysis

#### Preparation of grain mash samples for amino acids and peptide analysis

Grain mash samples (10 mL) were centrifuged at 13000 × g for 15 min at 20°C in a Hettich^®^ ROTANTA 460/460R centrifuge (Andreas Hettich GmbH & Co. KG, Tuttlingen, Germany) to separate the solids. The liquid phase was then filtered using Minisart RC 0.2μm syringe filters (Sartorius, Göttingen, Germany) to remove any residual biomass.

Sugars were removed from the grain mash by fermentation with a peptide transporter knockout (PepKO) strain ^[16, 19]^. The latter was necessary to avoid the loss of amino acids due to Maillard reactions when total amino acid analysis was carried out under high temperature and acidic conditions. First, the samples were heated at 75 °C for 20 min to remove any residual proteolytic activity. In addition, the uninoculated grain mash samples were freeze-dried in a Heto PowerDry PL3000 Freeze Dryer (Thermo Fisher Scientific, MA, USA) and reconstituted in MilliQ water to remove ethanol, which would otherwise prevent fermentation. To ensure proper fermentation with the PepKO stain, 100 μL of a nutrient stock (Table S2) containing minerals, vitamins, and 100 mg/L nitrogen from NH_4_^+^ was added to the samples. Because the PepKO strain used was a descendant of a wine stain, the nutrient stock was also supplied with 0.1% glucoamylase (MEGA PACIFIC TECHNOLOGY INC, Arcadia CA, USA) to break down maltose and maltotriose into more easily assimilable glucose ^[20]^. In addition, because the PepKO strain contained KanMX as a marker for the deletion of *FOT* peptide transporter genes ^[21]^, 200 μg/mL G418 (InvivoGen, Toulouse, France) was added to prevent microbial contamination. Fermentation with the PepKO strain was performed in 15 mL conical tubes (Eppendorf AG, Hamburg, Germany) at 32 °C and at 150 rpm. The lids of the tubes were slightly opened to prevent overpressure. The inoculum (20 μL) was prepared as previously described. The end of the fermentation process was indicated by the cells dropping from the culture medium. As the samples contained variable concentrations of residual sugars, the fermentation time was sample dependent. Cells were separated from the samples by centrifugation at 9600 × g for 10 min at 4 °C in a MicroCL 21R Microcentrifuge (Thermo Fisher Scientific, MA, USA) and filtered using Minisart RC 0.2 μm syringe filters (Sartorius, Göttingen, Germany).

The resulting samples were filtered to obtain a ≤ 2 kDa fraction using Vivaspin 2 centrifugal concentrators (Sartorius AG, Göttingen, Germany) according to the manufacturer’s instructions. The retentate was washed twice with 2 mL of MilliQ water. The permeate was freeze-dried and reconstituted in Milli-Q water to obtain the final sample.

#### Analysis of free and bound amino acids

Concentrations of free and peptide-bound amino acids were analysed on a Waters ACQUITY UPLC^®^ system (Waters Corporation, Milford, MA, USA) coupled to Waters ACQUITY UPLC tuneable ultraviolet (TUV) detector (Waters Corporation, Milford, MA, USA) after derivatisation using Waters AccQ-Tag chemistry, as described by Fiechter and Mayer ^[21]^. For total amino acids, the samples were first hydrolysed with 6M HCl + 1% (v/v) phenol at 105 °C for 22 h in vacuum using an Eldex H/D workstation (Eldex Laboratories Inc., Napa, CA, USA). The peptide-bound amino acids were calculated by subtracting the measured free amino acids from the total amino acids.

### Peptide analysis

#### Sample preparation

The SWM samples were first mixed (1:1) with methanol in BRANDplates™ pureGrade™ 96-Well Microplates (BRAND GMBH + CO KG, Wertheim, Germany). The plates were then centrifuged at 560 × g using a BioSan LMC-3000 centrifuge (Biosan, City, Company) to remove the precipitate. Then, 20 μL from each well was transferred to a Waters™ round well polypropylene 350 µL 96-well sample collection plate (Waters Corporation, Milford, MA, USA) and diluted with Milli-Q water (160 μL) to a final volume of 180 μL. All samples were spiked with 0.5 ppm caffeine (20 μL) to a final volume of 200 μL. Caffeine was used as an internal standard (reference housekeeping ion) during subsequent analysis.

Ice-cold grain mash samples (100 μL) were first mixed with 900 μL of ice-cold methanol in 1.5 mL Eppendorf^®^ tubes (Eppendorf, Hamburg, Germany) and centrifuged at 11,200 × g at 4 °C for 20 min in a MicroCL 21R Microcentrifuge (Thermo Fisher Scientific, MA, USA) to remove the precipitate. The samples were then freeze-dried in a Heto PowerDry PL3000 Freeze Dryer (Thermo Fisher Scientific, MA, USA) and reconstituted in 100 μL Milli-Q water. All samples were spiked with 0.5 ppm caffein to a final volume of 110 μL.

#### Liquid chromatography-mass spectrometry (ultra-high-pressure liquid chromatography ion mobility separation-enabled high-resolution mass spectrometry (UHPLC-IMS-HRMS))

The peptides were analysed using the methodology described by Arju et al. (2022). Briefly, the Waters I-Class Plus (SM-FL) UPLC system (Waters Corporation, Milford, MA, USA) was coupled with a Waters Vion IMS-QTof Mass Spectrometer equipped with a LockSpray II Exact Mass Source Enclosure and MKII tool-free ESI probe assembly directly connected to the column outlet. Nitrogen was used as collision gas. The instrument was controlled using Waters UNIFI 1.9.4 (3.1.0, Waters Corporation, Milford, MA, USA).

The instrument was operated in positive polarity, sensitivity mode (31,000 FWHM at 556.2766 m/z), and labile ion mobility tuning mode. The analysis type was set as Peptide Map (IMS), and the experiment type was set to MSe. Data were acquired in the HDMSe mode with a scan time of 0.165 s for synthetic and 0.250 s for grain mash. The following manual quadrupole profile was used for SWM: mass 150/250/450 (m/z), dwell time 60/20 (% scan time), ramp time 10/10 (% scan time); and for grain mash: mass 162/300/500 (m/z), dwell time 35/35 (% scan time), ramp time 15/15 (% scan time).

The injection volume was 5 µL. Full loop (5 µL) injection mode was used with a 3-volume overfill. The samples were analysed using an Acquity UPLC HSS T3 Column (1.8 µm, 1 × 150 mm, Waters Corporation, Milford, MA, USA) maintained at 45 °C. The initial flow rate was 0.15 mL/min. The gradient for synthetic mash was as follows: a 0–0.5 min hold at 1% B; 0.5-10.5 min linear gradient, 1-30% B; 10.5-12 min linear gradient, 30-95% B accompanied by a linear flow rate increase of 0.15-0.2 mL/min; 12-12.5 min hold at 95% B accompanied by a linear flow rate increase of 0.2-0.25 mL/min; 12.5-12.6 min linear gradient, 95-1% B accompanied by a linear flow rate decrease of 0.25-0.15 mL/min; and 12.6-15 min hold at 1% B. The gradient for grain mash was as follows: a 0-1 min gold at 1% B; 1-16 min linear gradient, 1-30% B; 16-17 min linear gradient, 30-95% B accompanied by a linear flow rate increase of 0.15-0.2 mL/min; a 17-17.5 min hold at 95% B accompanied with a linear flow rate increase of 0.2-0.3 mL/min; 17.5-17.6 min gradient, 95-1% B accompanied with linear flow rate decrease of 0.3-0.15 mL/min; a 17.6-20 min hold at 1% B.

## Data Treatment and Statistical Analysis

Data was treated and analysed using R v4.3.2 (R Core Team 2023) and RStudio (RStudio Team 2023). The growth rate and maximum cell population in SWM were estimated by fitting the optical density curves with the “summarizegrowth” R-function from the GrowthCurver package ^[22]^. The significance of ANOVA between the mean values was assessed using Tukey’s HSD method.

Peptide mapping for peptide analysis was conducted according to the method described by Arju et al. (2022) ^[23]^. The time window for peptide detection was extended to 14 min to accommodate the extended gradient used to analyse the grain mash samples. Peptide mapping in the grain mash samples of peptide masses was performed against five barley storage proteins and nine wheat storage proteins (Table S3).

Data processing of amino acids and peptides from the strain characterisation in SWM was performed, as described by Berg et al. ^[16]^. A list of the mass spectrometric data of di-to nonapeptides identified in the BSA hydrolysate in this study is shown in Table S6.

All data were visualised using ggplot2 ^[23]^.

## Data Availability

Data files and the R script for data treatment and analysis can be found at: https://zenodo.org/records/13118415

All data are contained in the manuscript and the supplemental material.

## Results

### Effect of oligopeptides on the fermentation kinetics in synthetic whisky mash

Fermentation experiments were performed on synthetic whisky mash (SWM) to study the uptake of peptides as a nitrogen source by the whisky yeast strains used in this study (Table 1) under different conditions of nitrogen supply (Table 2). In the first experiment, the fermentation medium (AP230) contained 130 mg N/L from free amino acids (FAA) and 100 mg N/L from peptides with a molecular weight (MW) ≤ 2 kDa. In the second medium (P230) the FAA were deficient (30 mg N/L), and peptides were the only substantial source of nitrogen (200 mg N/L). Finally, the effect of additional nitrogen supplementation (70 mg N/L) with either diammonium phosphate (DAP) or peptides was studied (media PP300 and PD300, respectively). The effect on the fermentation kinetics is shown in Figure 1.

**Figure 1.**
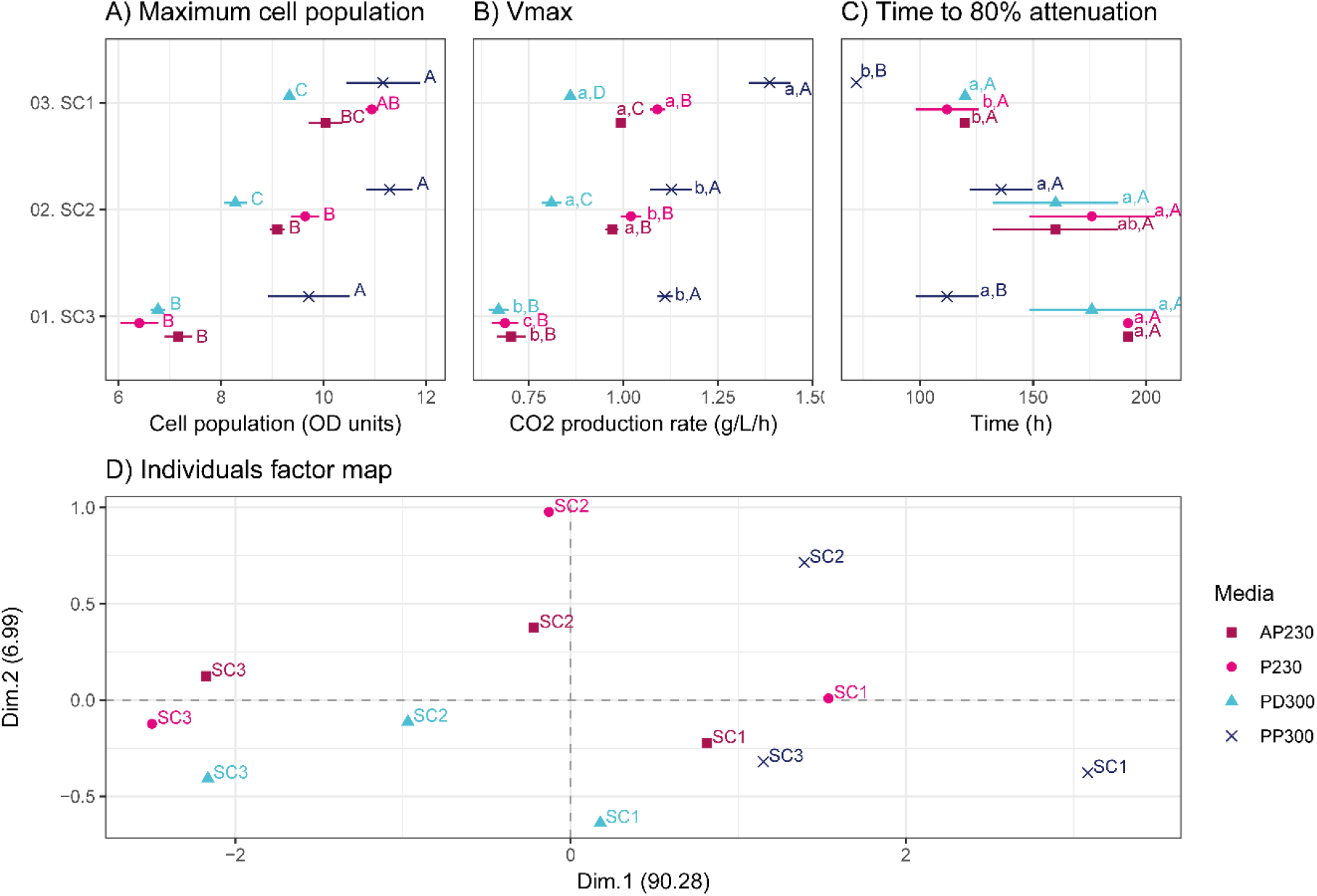
Cell growth and fermentation kinetics in synthetic whisky mash with different nitrogen conditions. Letters denote the statistical groups from Tukey’s tests (P < 0.05), with lower-case letters indicating different groups of strains per medium and upper-case letters indicating the groups of media for each strain. The “time to 80% attenuation” was chosen as not all fermentations reached 100% of attenuation during the set experimental time. Principal component analysis (PCA) (D) was performed using the three fermentation parameters depicted in A, B, and C.

In the experiments with a total N content of 230 mg N/L, containing either free amino acids (56.5% of N) and peptides(43.5% of N) (AP230) or only peptides (P230), the three strains displayed no significant differences in maximum cell population (Figure 1A), maximum CO_2_ production rate (Vmax, Figure 1B), and fermentation time (Figure 1C). Strain SC1 was the only strain that exhibited a significantly higher Vmax value in the P230 medium, resulting in a slight, yet statistically insignificant reduction in overall fermentation time (80% attenuation, determined based on the maximum theoretical CO_2_ production from 150 g/L maltose and glucose (9:1)). These findings corroborate previous results from the experiments we conducted with a wine yeast and suggest that peptides individually can also provide the necessary nitrogen to these whisky strains ^[16]^.

Among the three strains, strain SC3 exhibited the lowest fermentation performance (Vmax and fermentation time) at 230 mg N/L (Figure 1), which could be attributed to its higher nitrogen demand than that of the other strains. Interestingly, with additional nitrogen supplementation (70 mg N/L) with diammonium phosphate (DAP, medium PD300), the fermentation kinetics did not improve, whereas a positive impact was observed with an increased peptide concentration (medium PP300). Thus, in the PP300 medium, the maximum cell population and Vmax increased by 52% and 62%, respectively, while the fermentation time decreased by 42%, compared to the P230 medium. Similarly, strain SC1 in the PP300 medium showed a 28% increase in Vmax and a 36% decrease in fermentation time, although the biomass density remained similar to that in the P230 medium. In the case of strain SC2, the maximum cell population and Vmax increased by 17% and 11%, respectively, resulting in a 23% decrease in fermentation time, albeit not statistically different from the fermentation time in the lower nitrogen-containing P230 medium. Interestingly, DAP supplementation (PD300 medium) had a negative effect on the maximum cell population and Vmax by strain SC1 and SC2. Nevertheless, despite variations in the maximum cell population and Vmax, the fermentation time remained unaffected.

A principal component analysis plot was constructed to support these observations (Figure 1D). The experiments in the AP230 and P230 media clustered together by strain, indicating that peptides can provide the necessary nitrogen to these whisky strains when FAA are limited, without affecting fermentation efficiency. For strain SC3, the experiments conducted on the PD300 medium cluster with the AP230 and P230 lower nitrogen-containing media, showing no or minimal effect of ammonium supplementation on this strain. The negative effect of DAP supplementation on the other strains can be observed by their respective PD300 medium clustering away from their respective AP230 and P230 clusters. The positive effect of increased nitrogen content from peptide supplementation (PP300) on strain SC3 can be seen by its clustering with strain SC1 in the AP230 and P230 media. Strain SC3 in the PP300 medium also moved closer to this cluster, whereas strain SC1 in the PP300 medium formed its own group.

### Oligopeptide uptake by whisky yeast strains in synthetic whisky mash

In addition to the fermentation kinetics, peptide consumption by the three whisky yeast strains was monitored during the first 72 hours of fermentation (Figures 2, S1-4). Intensive uptake of peptides with different chain lengths (up to nine amino acids) simultaneously with FAA was observed in both AP230 and P230 media for all strains (Figure S4). The conserved ability of the three strains to take up peptides from the environment was also evident from their similar peptide consumption profiles in these two media.

**Figure 2.**
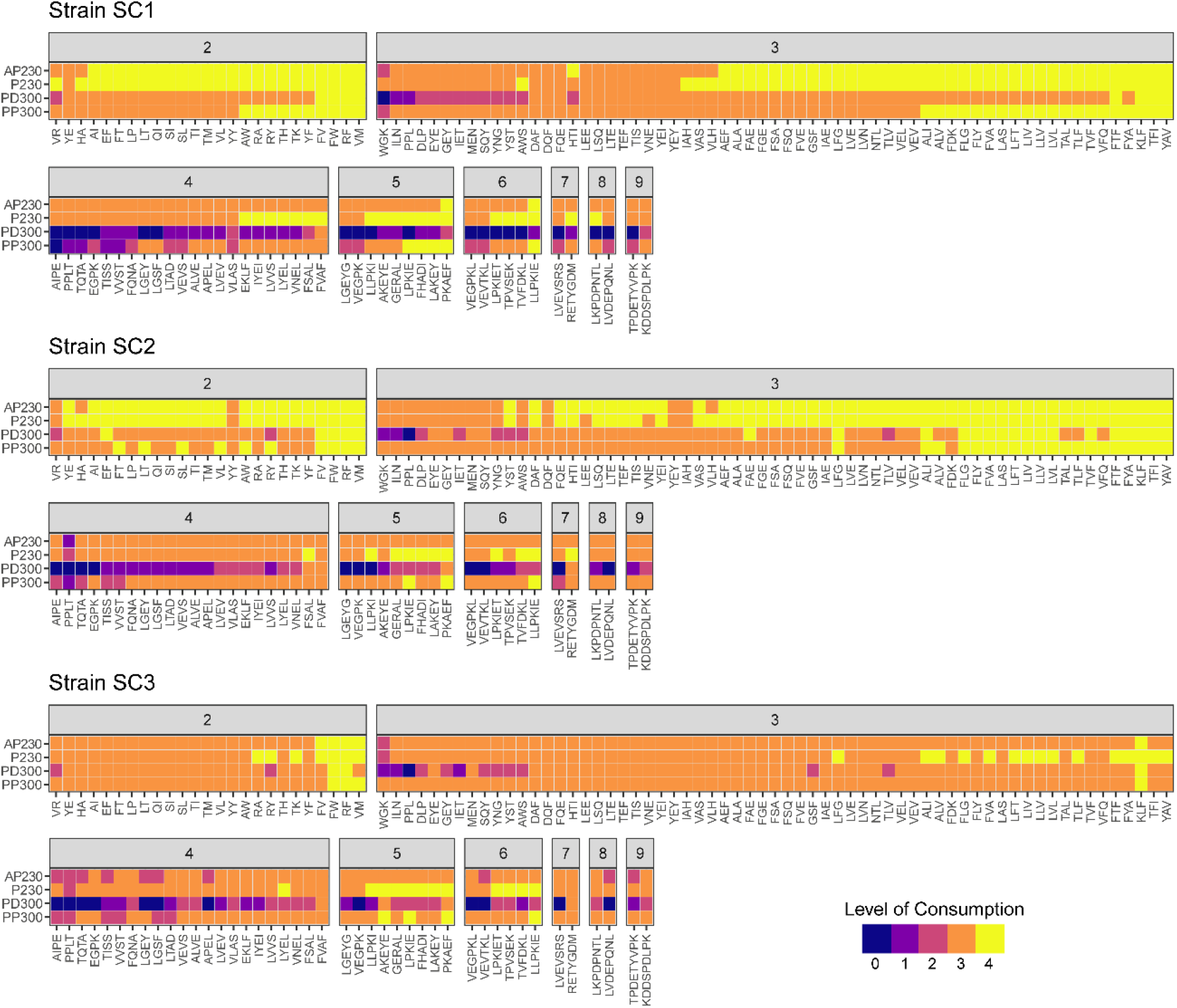
Consumption of peptides in synthetic whisky mash by the three whisky strains during the first 72 hours of fermentation. Peptides were grouped according to the number of amino acid residues: 2, dipeptides; 3, tripeptides; 4, tetrapeptides; 5, pentapeptides; 6, hexapeptides; 7, heptapeptides; 8, octapeptides ; and 9, nonapeptides. The area under the curve (AUC) was calculated from the relative abundance curve of each peptide. The AUC values were compared to those of a virtual negative control (100% abundance over 72 hours). The consumption of a particular peptide was level 0 when its abundance AUC was equal to or higher than 80% of the negative control AUC; level 1, 60–80%; level 2, 40–60%; level 3, 20–40%; and level 4, when the peptide abundance AUC was lower than 20% of the control AUC.

In the medium where additional nitrogen was added in the form of DAP (PD300), ammonia was consumed first, which resulted in delayed uptake of di- and tripeptides and limited the uptake of peptides with longer chain lengths in all strains (Figures 2, S1-4). In addition, the strains revealed distinct patterns of peptide uptake in the medium with increased peptide content (PP300), which may be attributed to their individual nitrogen requirements. For example, strain SC1 showed similar consumption levels for the di- and tripeptides in the PP300 medium as in the P230 medium (Figure 2), while peptides with a longer chain length were taken up more slowly. In strain SC2, this effect was less pronounced, whereas in strain SC3, all peptides were taken up simultaneously. As the methodology utilised in this study does not enable the exact quantification of individual peptides, it was impossible to determine the exact quantity of nitrogen contributed by each peptide length group to the overall nitrogen intake.

Together with the observed effects of peptides on fermentation performance (Figure 1), these results support our hypothesis that peptides can serve as a substantial alternative source of nitrogen for yeast.

### Nitrogen availability and the role of peptides in Scotch grain whisky fermentation

To study the role of peptides as a yeast nitrogen source in Scotch grain whisky fermentation, a grain mash sample from an industrial Scotch grain whisky production process was obtained, and fermentation was conducted in one-litre benchtop fermenters using the three strains previously characterised for their peptide uptake capability in SWM.

All strains completed fermentation in the grain mash, albeit with distinct fermentation kinetics (Figure 3). Strain SC1 showed the fastest fermentation, ceasing active fermentation by 24 hours. After this point, the CO_2_ production rate significantly decreased, and fermentation was complete after approximately 48 hours. In contrast, with strain SC2 and SC3 the CO_2_ production rate started to cease at 36 hours with fermentation completion at 48 hours. The ethanol content of the fermentation with strain SC2 was the highest at 8.8 ± 0.2 %(v/v), followed by strain SC3 at 8.7 ± 0.2 %(v/v), and strain SC1 at 8.5 ± 0.4 %(v/v), which were not significantly different from each other (α=0.05).

**Figure 3.**
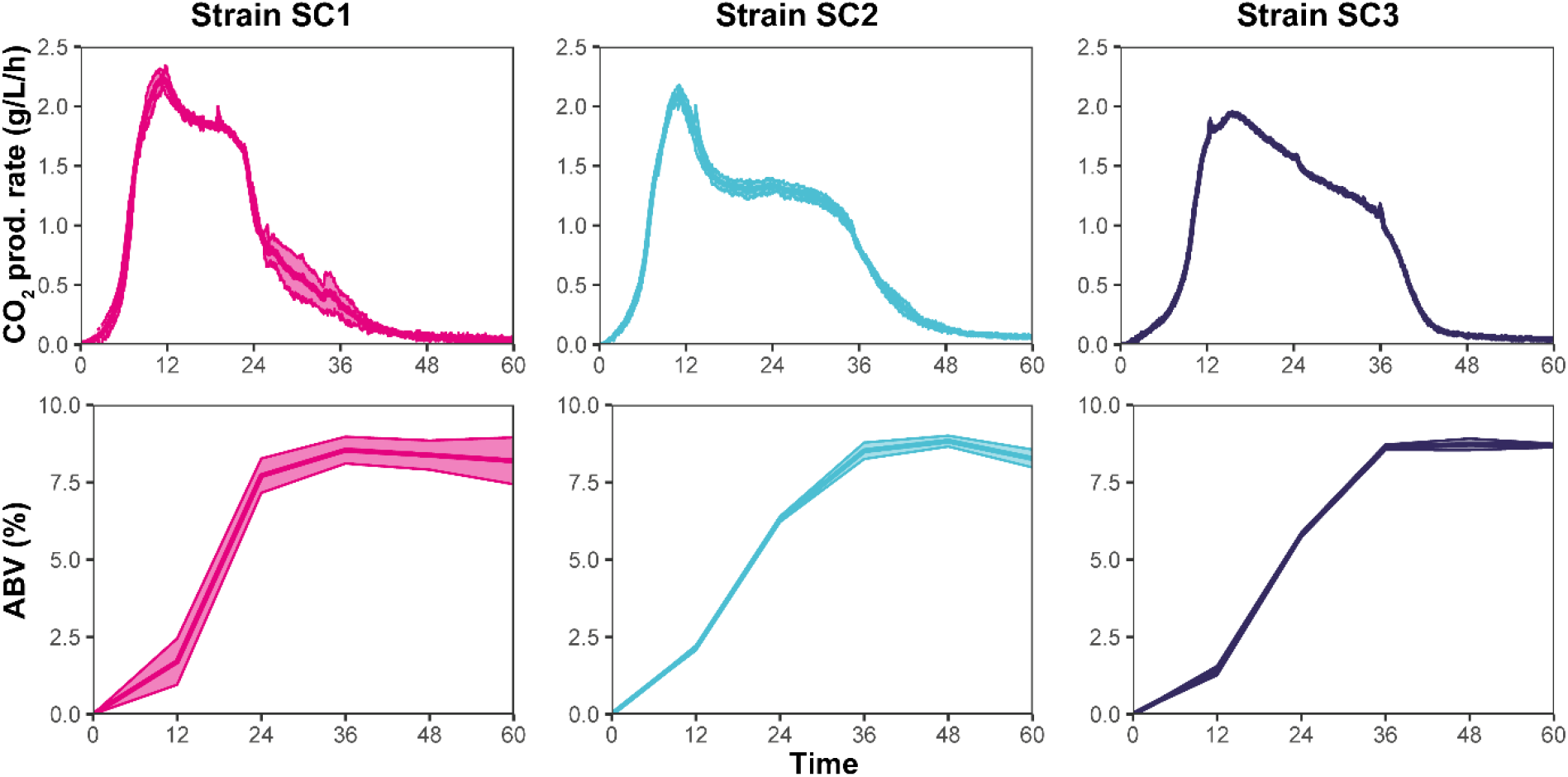
CO_2_ production rate and ethanol concentration by volume (ABV) during laboratory -scale simulated Scotch grain whisky fermentation with three commercial whisky strains.

The initial grain mash contained 90.2 ± 6.73 mg N/L from FAA (Figure 4). According to previously published studies, this amount does not meet the minimum requirement for satisfactory fermentation efficiency (150-250 mg N/L) for the given fermentable sugar content ^[5, 24]^. However, the grain mash also contained oligopeptides, some of which could be a potential source of N for yeast. In total, 70.4 ± 9.9 mg N/L was determined to be derived from peptides with a MW ≤ 2 kDa. Thus, the sampled grain mash initially contained up to 160.6 ± 12.0 mg N/L of potentially yeast-assimilable nitrogen.

**Figure 4.**
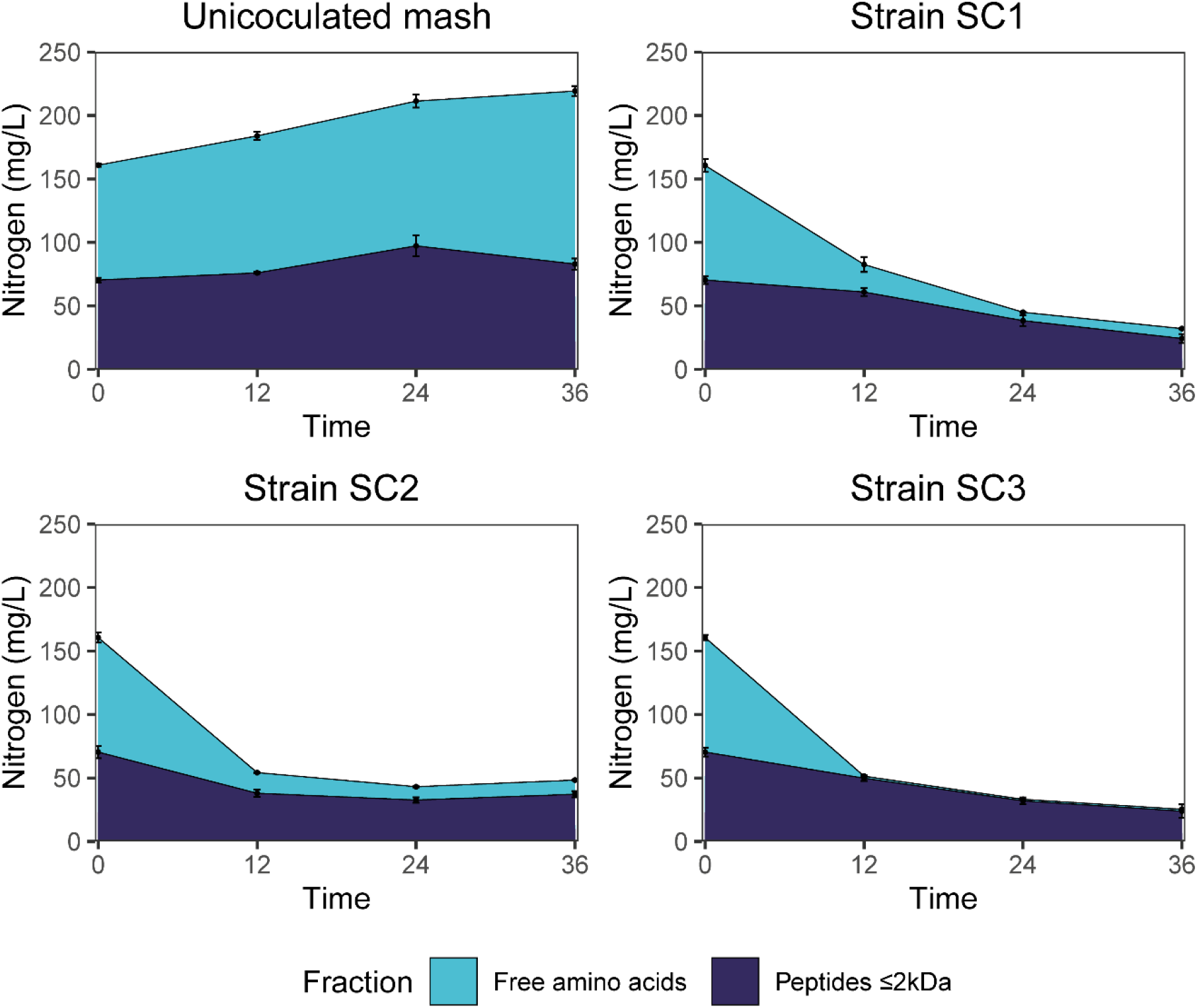
Changes in FAA and oligopeptide (≤ 2 kDa) content during fermentation of Scotch grain mash and uninoculated grain mash. One experiment with uninoculated grain mash, simulating the fermentation process with changing pH and EtOH concentration, was conducted to determine the background production of potentially available nitrogen sources for yeast (Table 3). Free proline was not taken into account as it is poorly assimilated by yeast during fermentation ^[4, 40]^.

The production of grain mash for Scotch grain whisky fermentation does not involve boiling, which could result in residual proteolytic activity during fermentation originating from the malt. To determine the amount of additional potentially yeast-assimilable nitrogen liberation, an experiment was conducted using an uninoculated grain mash, where the actual grain mash fermentation conditions were simulated by gradually changing the pH and ethanol concentration (Table 3, Figure 4). The results showed an increase in nitrogen content in both FAA and, to a lesser extent, peptides ≤ 2 kDa. The increase of potentially available nitrogen content amounted to a total of 58.4 ± 6.28 mg N/L in 36 hours, with 45.8 ± 4.1 mg N/L from FAA and 12.6 ± 4.7 mg N/L from peptides. Therefore, a total of 219.0 ± 13.5 mg N/L was available for fermentation, with 38% of it derived from peptides with a MW ≤ 2 kDa. Corroborating the observed fermentation kinetics and literature data, this quantity was sufficient to complete fermentation ^[5]^.

Although the strains displayed varying fermentation and nitrogen uptake kinetics, the amounts of nitrogen consumed from the grain mash were similar (Figure 4). Thus, strain SC1 consumed 186.8 ± 14.0 mg N/L, with 128.2 ± 7.9 mg N/L derived from FAA and 58.6 ± 11.5 mg N/L from peptides. Similarly, strain SC3 consumed 193.6 ± 14.5 mg N/L, with 134.5 ± 7.9 mg N/L derived from FAA and 59.1 ± 12.2 mg N/L from peptides. Strain SC2 consumed 175.3 ± 13.7 mg N/L, with 126.0 ± 7.9 mg N/L derived from FAA and 49.4 ± 11.2 mg N/L from peptides. Thus, peptide-derived nitrogen in these fermentations comprised ∼30% of the total nitrogen assimilated by yeast.

Peptide production, degradation and consumption (based on the change in their MS peak intensities) at different sampling points were compared (Figure 5). In total, 153 peptides ranging from 2 to 23 AA in length could be followed. Peptide annotations were obtained by mapping the peptide masses to the six most abundant barley storage proteins (hordeins) and nine wheat storage proteins (glutenins) (Table S3) ^[25, 26]^. However, this approach is subject to a degree of identification inaccuracy due to the numerous proteins that were included in peptide screening, as well as those that were not included. Consequently, we decided to attribute only the peptide length to the observed peptide candidate masses, as suggested in another study ^[26]^.

**Figure 5.**
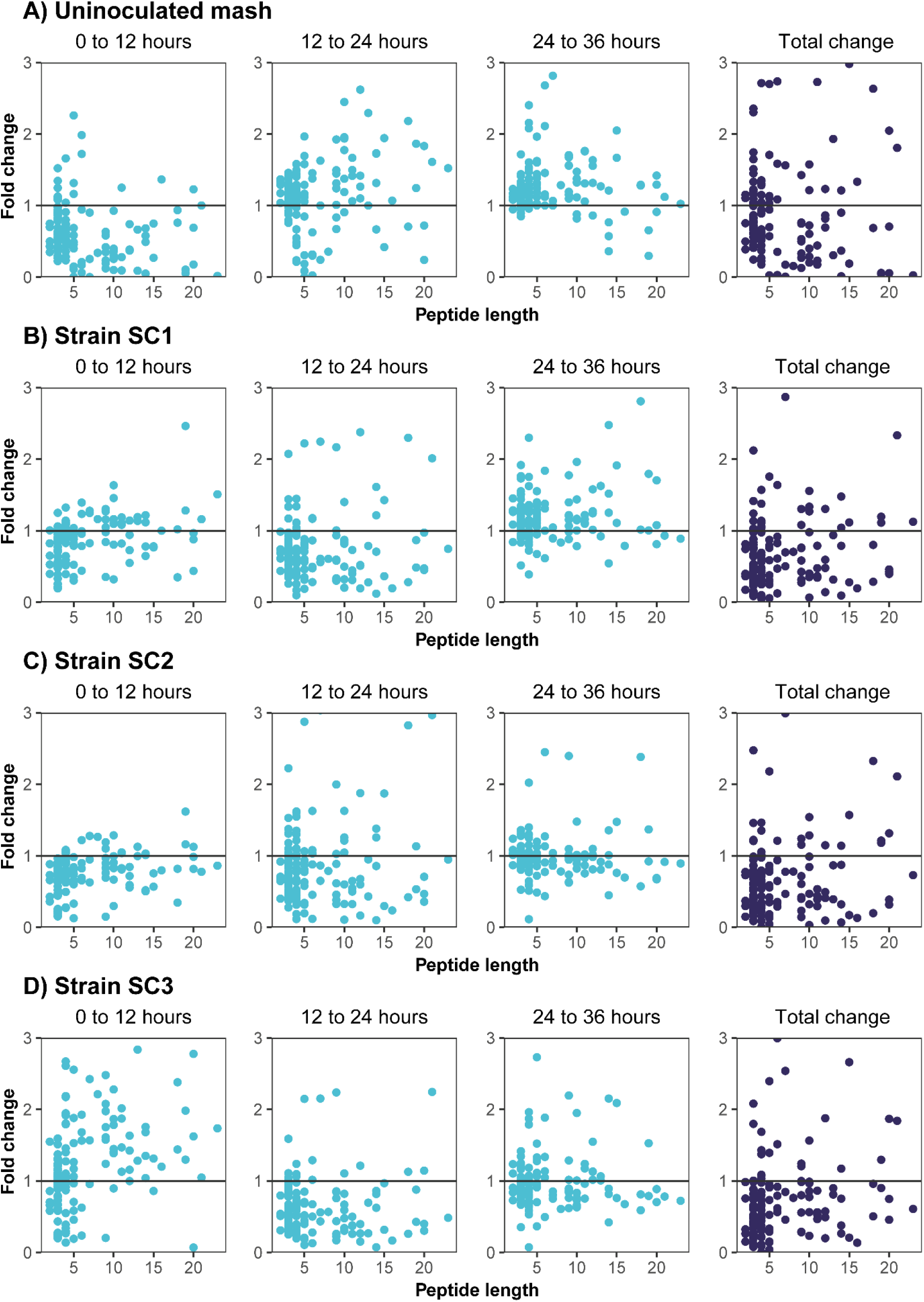
Fold changes in peptide abundance between samples of the uninoculated Scotch grain mash and the fermentations of the same mash with strains SC1-3. The “total change” depicts the fold change between peptides candidates at the 0-hour sample and the 36-hour sample.

Analysis of peptides in the uninoculated mash revealed complex dynamics of peptide production and degradation in the grain mash due to malt proteases (Figure 5A). During the first 12 hours of incubation, degradation of peptides occurred, as shown by the decreased relative concentration of most peptides, as well as the increased concentration of free amino acids (Figure 4). The hydrolysis continued between 12 and 36 hours, first with increasing relative concentrations of higher MW peptides (12 to 24 hours), which were then further degraded into smaller ones between 24 and 36 hours, suggesting an interplay between various proteases and peptidases. The simultaneous production and degradation of peptides also explains the limited change in the contribution of peptides to the overall increase in the potentially yeast-assimilable nitrogen content of the grain mash, as shown in Figure 4.

The complex kinetics of peptide formation and degradation due to proteolysis by malt-derived enzymes makes it difficult to precisely interpret the extent of their consumption by yeast (Figure 5B-D). Thus, the changes in the relative concentration of small MW peptides (two to five AA in chain length) during fermentation in the first 12 hours were very similar to those observed in the uninoculated mash incubation experiment. In contrast, a higher number of peptides with a negative fold change in the relative concentration during later stages of fermentation when free amino acids were depleted suggests their consumption by yeast. In all three strains, the net change between the start and end of fermentation suggested the consumption of di-to-pentapeptides (average fold change of 0.6 compared to 0.9 in the uninoculated mash).

The only major difference among the three yeast strains was observed in the experiment with strain SC3 (Figure 5D), in which the apparent increase in peptides was much more pronounced during the first stage of fermentation (0 to 12 hours). The exact reasons for this are unknown but may be related to the longer lag phase observed for this strain compared to the other two strains during which the consumption of peptides did not take place yet. Bound amino acid analysis of the 2 kDa fraction in strain SC3 did not show a substantial increase in the nitrogen content of peptides between 0 and 12 hours (Figure 4). Therefore, it was not possible to determine the significance of this observation without quantitative data on specific peptides at more frequent sampling points.

Overall, these findings indicate that the production, degradation, and consumption of peptides during Scotch grain whisky fermentation is a highly dynamic and complex process.

## Discussion

Around 90% of the raw material used in the production of Scotch grain whisky consists of unmalted grains, which may lead to nitrogen deficiency for yeast in the fermentable grain mash. This study investigated the availability and utilisation of nitrogen sources by yeast during the fermentation of Scotch grain whisky, with a particular focus on the role of oligopeptides. First, three commercial *Saccharomyces cerevisiae* whisky strains were assessed for their ability to take up peptides from peptide-supplemented synthetic whisky mash, using various nitrogen sources. All three strains were able to complete fermentation with peptides as the sole nitrogen source, demonstrating that peptides with a suitable chain length can satisfy the nitrogen requirements of yeast. We previously characterised a *Saccharomyces cerevisiae* wine yeast (59A, haploid variant of the commercial wine strain EC1118 ^[27]^) for its ability to take up peptides with different chain lengths in a similar analysis and found that it could utilise peptides up to seven amino acids in length, which were the highest chain length peptides determined by the analyses used ^[16]^. By further optimising the peptide mapping analysis, in the current study we suggest that the three whisky strains can take up peptides of up to at least nine amino acids in length. Like the wine strain 59A, these whisky strains took up smaller peptides first, with the only exception being that the whisky strains preferred di- and tripeptides, whereas in 59A, tetrapeptides were also preferred ^[16]^. This can be explained by the activity of the Fot family of peptide transporters, which is specific to wine yeasts ^[16, 19, 27, 28]^. Ptr2 (previously characterised as a di- and tripeptide transporter ^[29]^) is presumably the active di- and tripeptide transporter in whisky strains, explaining why tetrapeptides were taken up later with the other larger peptides.

The applied peptide mapping methodology in combination with using a single protein digest, allows for reliable peptide identification at the level of peptide length and amino acid composition but not the sequence ^[17]^. Peptides with different consumption levels shared similar chemical properties and could not be differentiated. Further investigation of the sequence-specific intake of peptides by these whisky strains necessitates experiments with synthetic peptides. Alternatively, proteases with known cleavage sites, such as trypsin or chymotrypsin, could be used to produce the BSA hydrolysate. By employing *in silico* methods, peptide mass data can then be utilised for peptide fingerprint identification.

Measurements taken from the Scotch grain mash obtained from an industrial distillery indicate that when a significant proportion of the mash bill is unmalted (approximately 90% in this case), the initial YAN content of the grain mash (∼160 mg N/L) is relatively low, considering that the minimum nitrogen requirement for satisfactory fermentation efficiency for the mash with the respective fermentable sugar concentration remains between 150 and 250 mg N/L ^[5]^ and that not all peptides in the ≤2 kDa molecular weight (MW) fraction might be fully assimilable by yeast. However, our study showed that the residual proteolytic activity that remained in the mash after conversion provided an additional 36% of potentially assimilable nitrogen to yeast during fermentation, of which the majority (80%) was in the form of free amino acids and 20% was contributed by peptides with MW ≤ 2 kDa. The fermentation experiments with three commercial whisky strains indicated that the final nitrogen content in the mash remained sufficient to support complete fermentation, with the strains taking up ∼ 185 mg N/L on average, of which ∼30% can be attributed to peptides. Thus, the remaining proteolytic activity in the mash seems essential for the fermentation process, and peptides form a considerable source of nitrogen for yeast during Scotch grain whisky fermentation.

Several protease families are involved in the germination process of barley, and are therefore present in mash ^[30]^. Heat-stable endoproteinases and carboxypeptidases may retain their activity in the mash during subsequent fermentation because the temperature of the conversion process does not exceed 70 °C ^[13, 14, 30]^, which was observed in our experiments. The LC-HRMS analysis of relative concentrations of individual peptides in uninoculated mash, in which the temperature, pH, and ethanol concentration were changed to simulate the fermentation process,suggested that the seconditions favoured the activity of both proteolytic enzyme classes of malt, resulting in the production of peptides and FAA. However, this process is dynamic and depends on the fermentation stage. The formation of free amino acids and small peptides occurred mostly at the beginning of fermentation (0 to 24 hours, T= 24 – 34 °C, pH= 5.2 – 4.3, EtOH= 0 – 50 g/L), whereas higher MW peptides started to accumulate in the later stages (12 hours and further). Because of this highly dynamic process, involving both production and degradation of peptides, limited variation in the nitrogen concentration in the MW ≤ 2 kDa peptide fraction of the uninoculated mash was observed. The extent of peptide uptake in the fermenting mash during the first phases of fermentation (0 to 12 hours) was difficult to estimate because of the co-occurring activity of different malt proteases; however, the consumption of di-to pentapeptides was more apparent between 12 and 36 hours.

The current MS-based methodology for studying the composition and concentration of peptides in unelucidated matrices, such as Scotch grain mash, has limitations. First, not all protein sequences in barley and wheat have been fully sequenced ^[31]^. Thus, in the present study, the obtained peptide masses were mapped only to the major (most abundant) known storage proteins in barley (hordeins) and wheat (glutenins) for annotation ^[25, 26]^. However, this approach is subject to a degree of identification inaccuracy due to the numerous protein sequences (total of 15) that were included for peptide mapping, as well as potentially unknown proteins that were not included but were also degraded into peptides. This limits the annotation of peptides to their size (number of AA residues in the chain length) instead of the exact AA sequence. As the MS peak intensity of peptides can be highly dependent on the amino acid composition and sequence ^[32]^, only the relative quantification of peptides by fold-change analysis of individual detected peptides was performed in this study. In combination with fractionation and subsequent A Aanalysis of the obtained molecular fractions, it was possible to explain these trends more quantitatively. However, some inaccuracies may have occurred in this case, for example due to retention of some smaller peptides on the cut-off filter. Ideally, the qualitative and quantitative analyses of peptides should be integrated into a single analysis. One method that currently exists makes use of the UV absorbance of peptides by coupling a photodiode array detector between LC and MS. The UV peak areas can be converted to absolute concentrations based on the law of Lambert-Beer, which requires the molar extinction coefficient of the peptide ^[33]^. However, this analysis would require extended chromatographical times to resolve most UV peaks in complex media, in addition to being hindered by the highly variable ionisation efficiencies of short peptides ^[33]^, which are required for the quantification of different peptides that co-elute under one UV peak ^[34]^. Further research to develop qualitative and quantitative analyses of peptides by MS in complex fermentation samples is therefore required.

Although the addition of nitrogen nutrients to grain whisky fermentation is not allowed by regulation in Scotland ^[34]^, it is allowed in other places. Therefore, the effects of additional nitrogen supplementation with either peptides or by the often industrially applied ammonium salt DAP on the fermentation performance in synthetic whisky mash were studied. In these experiments, supplementation with peptides resulted in higher biomass accumulation and CO_2_ production rates, and consequently, shorter fermentation times, than with the medium with a lower nitrogen content. This is in stark contrast to the effect of ammonium supplementation on fermentation performance, which was neutral or even negative. These results seem counterintuitive considering the popularity of DAP in industry; however, they are in line with what has been observed in other studies. In one study, nitrogen supplementation with a peptone induced higher biomass and ethanol production than ammonium sulphate ^[35]^. Another study revealed slower fermentation rates with ammonium compared to free amino acids, delivering equivalent amounts of nitrogen ^[36]^. Gene expression has been shown to be differentially reprogrammed based on the nitrogen source added ^[37]^: when ammonia was added, there was a higher expression of genes involved in amino acid biosynthesis, whereas the addition of amino acids resulted in a higher expression of genes related to protein biosynthesis. Thus, a possible explanation for the observed differences in the effect of nitrogen supplementation is that (peptide-derived) amino acids can be directly used in protein biosynthetic processes, while ammonium must first be incorporated into amino acids through *de novo* biosynthesis, requiring cellular resources including carbon and energy ^[38]^. Under the conditions used in this study, ammonia delayed peptide uptake, possibly through nitrogen catabolite repression ^[39]^. We hypothesised that this negative effect of ammonium supplementation on fermentation kinetics is due to the interplay between two factors: on the one hand, the cells need to *de novo* synthesise all the required amino acids, and on the other hand, the presence of ammonia hinders the uptake of peptides which could otherwise supply readily usable amino acids.

Strain SC3 is recommended for use in malted grain fermentation due to its high N-demand. However, our study suggests that this strain may also be suitable for grain fermentations with mostly unmalted grains, albeit with lower efficiency than Strain SC1. To achieve optimal process efficiency, it is essential to consider the available nitrogen content of the mash and the method of nitrogen supply to the yeast. This was demonstrated by our results on synthetic whisky mash, where strain SC3 greatly benefitted from additional nitrogen supplementation in form of small peptides. In the production of Scotch grain whisky, the use of malts with higher proteolytic strength or increasing the proportion of malted to unmalted grains can accommodate a specific yeast strain with a higher nitrogen demand. Outside of Scotland, it is permissible to use exogenous proteases to liberate free amino acids and peptides from the protein fraction of the mash or to add external nitrogen sources to the process. Increasing the nitrogen content in grain fermentation with the appropriate source of nitrogen may open the door to other strains that can be applied in this process.

In conclusion, these results show that peptides can play a considerable role as nitrogen sources to yeast in Scotch grain whisky fermentation. The three whisky yeast strains were able to take up a wide variety of peptides and use them effectively to attain complete fermentation. In industrial Scotch grain mash, the concentration of yeast assimilable nitrogen, including that produced during fermentation owing to residual proteolytic activity proved to be sufficient, with peptides being a considerable source of nitrogen.

## Acknowledgements

The authors want to acknowledge Martin Johannes Talu for his assistance with the experiments, Struan Reid for his advice throughout the project, and Mairéad Walsh for her help in editing this document.

## Disclosure statement

All authors have read and agreed to the published version of the manuscript and declare that the research was conducted in the absence of any commercial or financial relationships that could be construed as a potential conflict of interest.

## Funding

This work was funded by the Estonian Research Council project RESTA13.

## Notes

https://zenodo.org/records/13118415

### Competing Interest Statement

The authors have declared no competing interest.

### Summary of Updates

The referencing style was changed, extra discussion regarding method limitations and industy relevant outcomes of this article were added, strains were anonymised.

