## Supplemental Tables S1-7, Supplemental Figures S1-5 for "Nitrogen Availability and Utilisation of Oligopeptides by Yeast in Industrial Scotch Grain Whisky Fermentation"

### Supplemental Materials

**Table S1 – Amino acid concentrations used in the synthetic whisky mash (SWM).** Total "yeast-assimilable nitrogen" was 115 mg/L. Free proline was not taken into account as it is not or poorly assimilated by yeasts during fermentation.

| | Concentration (mg/L) | N (mg/L) | % total N | | Concentration (mg/L) | N (mg/L) | % total N |
|---|---|---|---|---|---|---|---|
| **Trp** | 45.47 | 6.24 | 4.76 | **Ala** | 50.48 | 7.94 | 6.06 |
| **Phe** | 54.12 | 4.59 | 3.50 | **Thr** | 25.15 | 2.96 | 2.26 |
| **Leu** | 72.13 | 7.70 | 5.88 | **Glu** | 50.28 | 4.79 | 3.65 |
| **Ile** | 27.16 | 2.90 | 2.21 | **Asp** | 47.15 | 5.00 | 3.82 |
| **Val** | 45.60 | 5.45 | 4.16 | **Gly** | 17.77 | 3.32 | 2.53 |
| **Met** | 15.36 | 1.44 | 1.10 | **Arg** | 59.53 | 19.15 | 14.62 |
| **Tyr** | 27.06 | 2.09 | 1.60 | **Gln** | 66.64 | 12.77 | 9.75 |
| **Lys** | 47.58 | 9.12 | 6.96 | **Ser** | 27.92 | 3.72 | 2.84 |
| **Cys** | 5.56 | 0.64 | 0.49 | **Asn** | 42.12 | 8.93 | 6.82 |
| **Pro** | 130.95 | 15.93 | 12.16 | **His** | 23.29 | 6.31 | 4.82 |

**Table S2 – Composition of the stock solution used in the removal of sugars from grain mash samples using the PepKO strain.** Components were added to supply already fermented samples with enough nutrients for the PepKO strain to remove sugars through refermentation. The concentrations are based on a recipe for synthetic grape must reported by Salmon and Barre (1998). The enzyme concentration was set as 0.1% w/w..

| **Component** | **Sample** mg/L | **PepKO Stock** mg/100mL | **Manufacturer** |
|---|---|---|---|
| $KH_2PO_4$ | 750 | 825 | Merck KGaA, Darmstadt, Germany |
| $K_2SO_4$ | 500 | 550 | Merck KGaA, Darmstadt, Germany |
| $MgSO_4 \cdot 7H_2O$ | 250 | 275 | Thermo Fisher Scientific Inc., Waltham, MA, USA |
| $CaCl_2 \cdot 2H_2O$ | 155 | 170.5 | Thermo Fisher Scientific Inc., Waltham, MA, USA |
| $(NH_4)_2PO_4$ | 471 | 518.1 | Thermo Fisher Scientific Inc., Waltham, MA, USA |
| $MnSO_4 \cdot H_2O$ | 4 | 4.4 | Thermo Fisher Scientific Inc., Waltham, MA, USA |
| $ZnSO_4$ | 4 | 4.4 | Thermo Fisher Scientific Inc., Waltham, MA, USA |
| $CuSO_4 \cdot 5H_2O$ | 1 | 1.1 | Merck KGaA, Darmstadt, Germany |
| KI | 1 | 1.1 | Thermo Fisher Scientific Inc., Waltham, MA, USA |
| $CoCl_2 \cdot 6H_2O$ | 0.4 | 0.44 | Merck KGaA, Darmstadt, Germany |
| $H_3BO_3$ | 1 | 1.1 | Thermo Fisher Scientific Inc., Waltham, MA, USA |
| $NaMoO_4 \cdot 2H_2O$ | 1 | 1.1 | Thermo Fisher Scientific Inc., Waltham, MA, USA |
| Myoinositol | 20 | 22 | Biosynth, Staad, Switzerland |
| Nicotinic acid | 2 | 2.2 | Thermo Fisher Scientific Inc., Waltham, MA, USA |
| Ca-pantothenate | 1.5 | 1.65 | Thermo Fisher Scientific Inc., Waltham, MA, USA |
| Thiamin-HCl | 0.25 | 0.275 | Merck KGaA, Darmstadt, Germany |
| Pyridoxine-HCl | 0.25 | 0.275 | Cayman Chemical Company, Ann Arbor, MI, USA |
| Biotin | 0.003 | 0.0033 | Merck KGaA, Darmstadt, Germany |
| Glucoamylase | 1 (g) | 1.1 (g) | MEGA PACIFIC TECHNOLOGY INC, Arcadia CA, USA |

**Table S3 - Hordein and glutenin UniProt entries used in peptide mapping of the grain mash.**

| Hordeins | | | | | |
|---|---|---|---|---|---|
| **Protein name** | **Entry name** | **UniProt Entry** | **Protein name** | **Entry name** | **UniProt Entry** |
| B1-hordein | HOR1_HORVU | P06470 | Gamma-hordein 1 | HOG1_HORVU | P17990 |
| B3-hordein | HOR3_HORVU | P06471 | Gamma-hordein 1 | HOG3_HORVU | P80198 |
| C-hordein | HOR7_HORVU | P06472 | | | |

| Glutenins | | | | | |
|---|---|---|---|---|---|
| **Protein name** | **Entry name** | **UniProt Entry** | **Protein name** | **Entry name** | **UniProt Entry** |
| Glutenin low MW subunit | GLTA_WHEAT | P10385 | Glutenin low MW subunit PC237 | GLT2_WHEAT | P02862 |
| Glutenin low MW subunit 1D1 | GLTB_WHEAT | P10386 | Glutenin low MW subunit 12 | GLT3_WHEAT | P08488 |
| Glutenin low MW subunit PTDUCD1 | GLTC_WHEAT | P16315 | Glutenin low MW subunit PW212 | GLT4 WHEAT | P08489 |
| Glutenin low MW subunit DY10 | GLT0_WHEAT | P10387 | Glutenin low MW subunit DX5 | GLT5_WHEAT | P10388 |
| Glutenin low MW subunit PC256 | GLT1_WHEAT | P02861 | | | |

**Tables S4-6 – Mean values of growth and fermentation parameters within the different media.** An analysis of variance followed by a Tukey's test (p-value = 0.05) was performed to indicate the significant differences among the values, which are represented in the "groups" column. Significant differences among the strains within the same condition are indicated with low case letters, and significant differences between conditions for the same strain are indicated with upper case letters. 'SD': standard deviation.

**Table S4 – Maximum population (k).** The k is presented in OD units. Significant differences between media were not assessed as yeast strains might have different cell sizes.

| | Strain SC1 | | | Strain SC2 | | | Strain SC3 | | |
|---|---|---|---|---|---|---|---|---|---|
| **Medium** | **k** | **SD** | **Groups** | **k** | **SD** | **Groups** | **k** | **SD** | **Groups** |
| AP230 | 10.04 | 0.33 | BC | 9.10 | 0.14 | B | 7.16 | 0.26 | B |
| P230 | 10.94 | 0.12 | AB | 9.64 | 0.27 | B | 6.40 | 0.37 | B |
| PD300 | 9.33 | 0.04 | C | 8.28 | 0.23 | C | 6.77 | 0.14 | B |
| PP300 | 11.16 | 0.72 | A | 11.29 | 0.45 | A | 9.71 | 0.80 | A |

**Table S5 – Maximum rate of CO2 production (Vmax).** The Vmax is presented in g/L/h.

| | Strain SC1 | | | Strain SC2 | | | Strain SC3 | | |
|---|---|---|---|---|---|---|---|---|---|
| **Medium** | **Vmax** | **SD** | **Groups** | **Vmax** | **SD** | **Groups** | **Vmax** | **SD** | **Groups** |
| AP230 | 0.99 | 0.01 | a,C | 0.97 | 0.02 | a,B | 0.70 | 0.04 | b,B |
| P230 | 1.09 | 0.02 | a,B | 1.02 | 0.03 | b,B | 0.69 | 0.04 | c,B |
| PD300 | 0.86 | 0.01 | a,D | 0.81 | 0.03 | a,C | 0.67 | 0.03 | b.B |
| PP300 | 1.39 | 0.06 | a,A | 1.13 | 0.06 | b,A | 1.11 | 0.02 | b,A |

**Table S6 – Time to reach 80% of attenuation (t80).** The t80 is presented in hours.

| | Strain SC1 | | | Strain SC2 | | | Strain SC3 | | |
|---|---|---|---|---|---|---|---|---|---|
| **Medium** | **t80** | **SD** | **Groups** | **t80** | **SD** | **Groups** | **t80** | **SD** | **Groups** |
| AP230 | 120 | 0 | b,A | 160 | 28 | ab,A | 192 | 0 | a,A |
| P230 | 112 | 14 | b,A | 176 | 28 | a,A | 192 | 0 | a,A |
| PD300 | 120 | 0 | a,A | 160 | 28 | a,A | 176 | 28 | a,A |
| PP300 | 72 | 0 | b,B | 136 | 14 | a,A | 112 | 14 | a,B |

**Table S7 – Mass spectrometric data of all di-nonapeptide candidates followed in the synthetic whisky mash.** RT: retention time; m/z: mass to charge ratio; z: charge, CCS: collisional cross-section; RSD: relative standard deviation (calculated from each fermentation starting point).

| Peptide ID | RT (min) | m/z | z | CCS (Å$^2$) | RSD (%) |
|---|---|---|---|---|---|
| AI | 3.33 | 203.1389 | 1 | 154.98 | 3.22 |
| AW | 5.13 | 276.1343 | 1 | 161.41 | 1.93 |
| EF | 4.39 | 295.1286 | 1 | 169.26 | 2.18 |
| FT | 2.92 | 267.1339 | 1 | 165.25 | 1.95 |
| FV | 5.02 | 265.1546 | 1 | 167.07 | 2.48 |
| FW | 7.99 | 352.1656 | 1 | 183.07 | 2.03 |
| HA | 1.43 | 227.1138 | 1 | 150.20 | 2.25 |
| LP | 4.42 | 229.1545 | 1 | 155.14 | 2.89 |
| LT | 2.03 | 233.1496 | 1 | 158.33 | 2.15 |
| QI | 3.29 | 260.1603 | 1 | 165.57 | 3.41 |
| RA | 1.89 | 246.1560 | 1 | 157.66 | 5.13 |
| RF | 2.89 | 322.1873 | 1 | 177.00 | 2.47 |
| RY | 1.54 | 338.1822 | 1 | 178.20 | 2.12 |
| SI | 1.02 | 219.1338 | 1 | 154.01 | 2.93 |
| SL | 3.10 | 219.1339 | 1 | 155.70 | 2.02 |
| TH | 2.58 | 257.1241 | 1 | 158.84 | 2.19 |
| TI | 3.44 | 233.1496 | 1 | 158.33 | 1.95 |
| TK | 2.57 | 248.1604 | 1 | 157.56 | 2.22 |
| TM | 2.07 | 251.1059 | 1 | 155.73 | 2.61 |
| VL | 4.57 | 231.1704 | 1 | 161.87 | 2.13 |
| VM | 3.32 | 249.1264 | 1 | 159.22 | 4.34 |
| VR | 2.60 | 274.1873 | 1 | 166.66 | 2.27 |
| YE | 2.16 | 311.1237 | 1 | 175.65 | 2.15 |
| YF | 6.03 | 329.1495 | 1 | 183.89 | 2.05 |
| YY | 4.17 | 345.1446 | 1 | 186.92 | 2.22 |
| AEF | 5.11 | 366.1660 | 1 | 184.41 | 2.17 |
| ALA | 3.59 | 274.1764 | 1 | 168.40 | 2.62 |
| ALI | 7.28 | 316.2230 | 1 | 180.79 | 2.47 |
| ALV | 5.64 | 302.2071 | 1 | 176.00 | 3.91 |
| AWS | 4.28 | 363.1662 | 1 | 184.51 | 2.67 |
| DAF | 5.01 | 352.1502 | 1 | 177.73 | 2.82 |
| DLP | 3.55 | 344.1811 | 1 | 181.56 | 3.35 |
| DQF | 4.61 | 409.1715 | 1 | 197.66 | 2.82 |
| EYE | 2.98 | 440.1661 | 1 | 200.48 | 4.07 |
| FAE | 4.11 | 366.1657 | 1 | 184.41 | 2.58 |

| Peptide ID | RT (min) | m/z | z | CCS (Å$^2$) | RSD (%) |
|---|---|---|---|---|---|
| FDK | 2.77 | 205.1076 | 2 | 252.55 | 1.85 |
| FGE | 4.03 | 352.1500 | 1 | 179.50 | 2.72 |
| FLG | 5.94 | 336.1915 | 1 | 185.43 | 3.61 |
| FLY | 7.94 | 442.2333 | 1 | 206.02 | 2.58 |
| FQE | 4.01 | 423.1872 | 1 | 200.95 | 3.03 |
| FSA | 4.55 | 324.1552 | 1 | 175.16 | 1.70 |
| FSQ | 3.20 | 381.1767 | 1 | 189.35 | 2.27 |
| FTF | 8.56 | 414.2023 | 1 | 197.51 | 2.43 |
| FVA | 5.27 | 336.1915 | 1 | 181.84 | 3.19 |
| FVE | 4.84 | 394.1973 | 1 | 194.42 | 2.49 |
| FYA | 8.45 | 400.1865 | 1 | 192.41 | 2.59 |
| GEY | 3.20 | 368.1452 | 1 | 180.77 | 2.53 |
| GSF | 4.06 | 310.1396 | 1 | 168.68 | 2.61 |
| HTI | 2.89 | 370.2081 | 1 | 191.52 | 2.98 |
| IAE | 3.11 | 332.1816 | 1 | 176.64 | 1.98 |
| IAH | 1.40 | 340.1977 | 1 | 183.49 | 2.34 |
| IET | 3.55 | 362.1921 | 1 | 188.15 | 2.23 |
| ILN | 4.19 | 359.2288 | 1 | 188.25 | 2.55 |
| KLF | 6.24 | 204.1360 | 2 | 252.62 | 2.44 |
| LAS | 3.70 | 290.1707 | 1 | 169.46 | 4.41 |
| LEE | 3.38 | 390.1871 | 1 | 190.89 | 2.32 |
| LFG | 7.53 | 336.1915 | 1 | 178.27 | 2.88 |
| LFT | 6.41 | 380.2178 | 1 | 193.03 | 2.02 |
| LIV | 7.42 | 344.2542 | 1 | 188.76 | 2.54 |
| LLV | 7.94 | 344.2545 | 1 | 188.76 | 2.28 |
| LSQ | 2.42 | 347.1924 | 1 | 181.45 | 2.15 |
| LTE | 3.27 | 362.1922 | 1 | 182.74 | 1.93 |
| LVE | 4.06 | 360.2130 | 1 | 184.60 | 2.00 |
| LVL | 7.69 | 344.2543 | 1 | 190.58 | 2.29 |
| LVN | 3.19 | 345.2136 | 1 | 181.52 | 2.42 |
| MEN | 1.42 | 393.1436 | 1 | 188.99 | 2.48 |
| NTL | 3.69 | 347.1923 | 1 | 188.66 | 3.44 |
| PPL | 5.33 | 326.2072 | 1 | 178.63 | 3.01 |
| SQY | 3.04 | 397.1717 | 1 | 188.87 | 1.73 |
| TAL | 4.37 | 304.1864 | 1 | 175.92 | 3.13 |
| TEF | 5.14 | 396.1763 | 1 | 192.53 | 2.27 |
| TFI | 8.25 | 380.2179 | 1 | 194.86 | 2.56 |
| TIS | 2.51 | 320.1815 | 1 | 175.31 | 2.76 |
| TLF | 7.74 | 380.2178 | 1 | 189.38 | 2.90 |
| TLV | 4.16 | 332.2178 | 1 | 181.98 | 2.84 |
| TVF | 6.38 | 366.2021 | 1 | 182.61 | 3.02 |
| VAS | 2.24 | 276.1558 | 1 | 164.85 | 3.44 |
| VEL | 6.11 | 360.2128 | 1 | 191.85 | 2.71 |

| Peptide ID | RT (min) | m/z | z | CCS (Å$^2$) | RSD (%) |
|---|---|---|---|---|---|
| VEV | 4.74 | 346.1967 | 1 | 186.88 | 5.02 |
| VFQ | 4.33 | 393.2129 | 1 | 192.62 | 2.50 |
| WGK | 5.84 | 390.2131 | 1 | 189.07 | 3.28 |
| VLH | 2.72 | 184.6178 | 2 | 248.00 | 3.64 |
| VNE | 1.91 | 361.1716 | 1 | 180.99 | 1.96 |
| YAV | 6.34 | 352.1866 | 1 | 179.50 | 2.44 |
| YEI | 6.61 | 424.2076 | 1 | 210.29 | 3.33 |
| YEY | 5.14 | 474.1870 | 1 | 210.82 | 2.43 |
| YNG | 1.70 | 353.1453 | 1 | 184.84 | 2.40 |
| YST | 3.01 | 370.1606 | 1 | 187.89 | 3.22 |
| AIPE | 4.70 | 429.234 | 1 | 198.93 | 3.26 |
| ALVE | 5.25 | 431.2499 | 1 | 202.58 | 2.42 |
| APEL | 5.84 | 429.2340 | 1 | 202.64 | 2.73 |
| EGPK | 2.74 | 430.2294 | 1 | 195.22 | 3.49 |
| EKLF | 8.79 | 536.3076 | 1 | 226.55 | 3.46 |
| FQNA | 3.61 | 479.2247 | 1 | 214.48 | 3.17 |
| FSAL | 7.33 | 437.2391 | 1 | 206.15 | 3.23 |
| FVAF | 9.07 | 483.2601 | 1 | 212.48 | 3.25 |
| IYEI | 8.14 | 537.2917 | 1 | 230.43 | 2.93 |
| LGEY | 5.31 | 481.2292 | 1 | 206.87 | 2.21 |
| LGSF | 6.70 | 423.2239 | 1 | 199.10 | 2.69 |
| LTAD | 3.28 | 419.2134 | 1 | 195.53 | 3.34 |
| LVEV | 6.67 | 459.2809 | 1 | 211.20 | 3.53 |
| LVVS | 5.42 | 417.2705 | 1 | 202.98 | 2.62 |
| LYEL | 6.92 | 537.2916 | 1 | 228.47 | 3.81 |
| PPLT | 4.85 | 427.2547 | 1 | 202.70 | 3.09 |
| TISS | 2.19 | 407.2129 | 1 | 195.87 | 2.12 |
| TQTA | 1.37 | 420.2086 | 1 | 199.18 | 3.50 |
| VEVS | 3.65 | 433.2292 | 1 | 200.67 | 2.31 |
| VLAS | 4.41 | 389.2391 | 1 | 192.74 | 3.79 |
| VNEL | 5.23 | 474.2556 | 1 | 214.61 | 3.58 |
| VVST | 3.16 | 405.2344 | 1 | 194.09 | 2.12 |
| AKEYE | 2.78 | 320.1521 | 2 | 280.41 | 3.47 |
| FHADI | 5.11 | 301.6495 | 2 | 281.28 | 3.62 |
| GERAL | 3.77 | 273.1556 | 2 | 276.39 | 2.47 |
| LAKEY | 4.12 | 312.1724 | 2 | 280.78 | 3.06 |
| LGEYG | 5.04 | 538.2506 | 1 | 218.79 | 3.31 |
| LLPKI | 7.14 | 292.2121 | 2 | 291.39 | 2.58 |
| LPKIE | 6.13 | 300.1913 | 2 | 287.76 | 2.58 |
| PKAEF | 4.47 | 296.1597 | 2 | 287.96 | 2.31 |
| VEGPK | 2.74 | 265.2988 | 2 | 273.65 | 3.23 |
| LLPKIE | 7.36 | 356.7333 | 2 | 317.94 | 2.58 |
| LPKIET | 6.48 | 350.7151 | 2 | 295.13 | 2.48 |

| Peptide ID | RT (min) | m/z | z | CCS (Å$^2$) | RSD (%) |
|---|---|---|---|---|---|
| TPVSEK | 2.93 | 330.6806 | 2 | 292.77 | 3.46 |
| TVFDKL | 7.79 | 361.7073 | 2 | 297.92 | 2.51 |
| VEGPKL | 5.87 | 321.6944 | 2 | 283.53 | 2.32 |
| VEVTKL | 6.22 | 344.7151 | 2 | 298.65 | 2.66 |
| LPKIETM | 7.53 | 416.2352 | 2 | 312.22 | 3.01 |
| LVEVSRS | 4.74 | 395.2264 | 2 | 306.41 | 3.13 |
| RETYGDM | 4.93 | 436.1833 | 2 | 311.50 | 2.42 |
| LKPDPNTL | 6.15 | 449.2549 | 2 | 317.69 | 3.41 |
| LVDEPQNL | 7.14 | 464.2414 | 2 | 320.52 | 3.57 |
| KDDSPDLPK | 4.71 | 507.7574 | 2 | 332.59 | 3.50 |
| TPDETYVPK | 5.58 | 525.2604 | 2 | 356.14 | 5.18 |

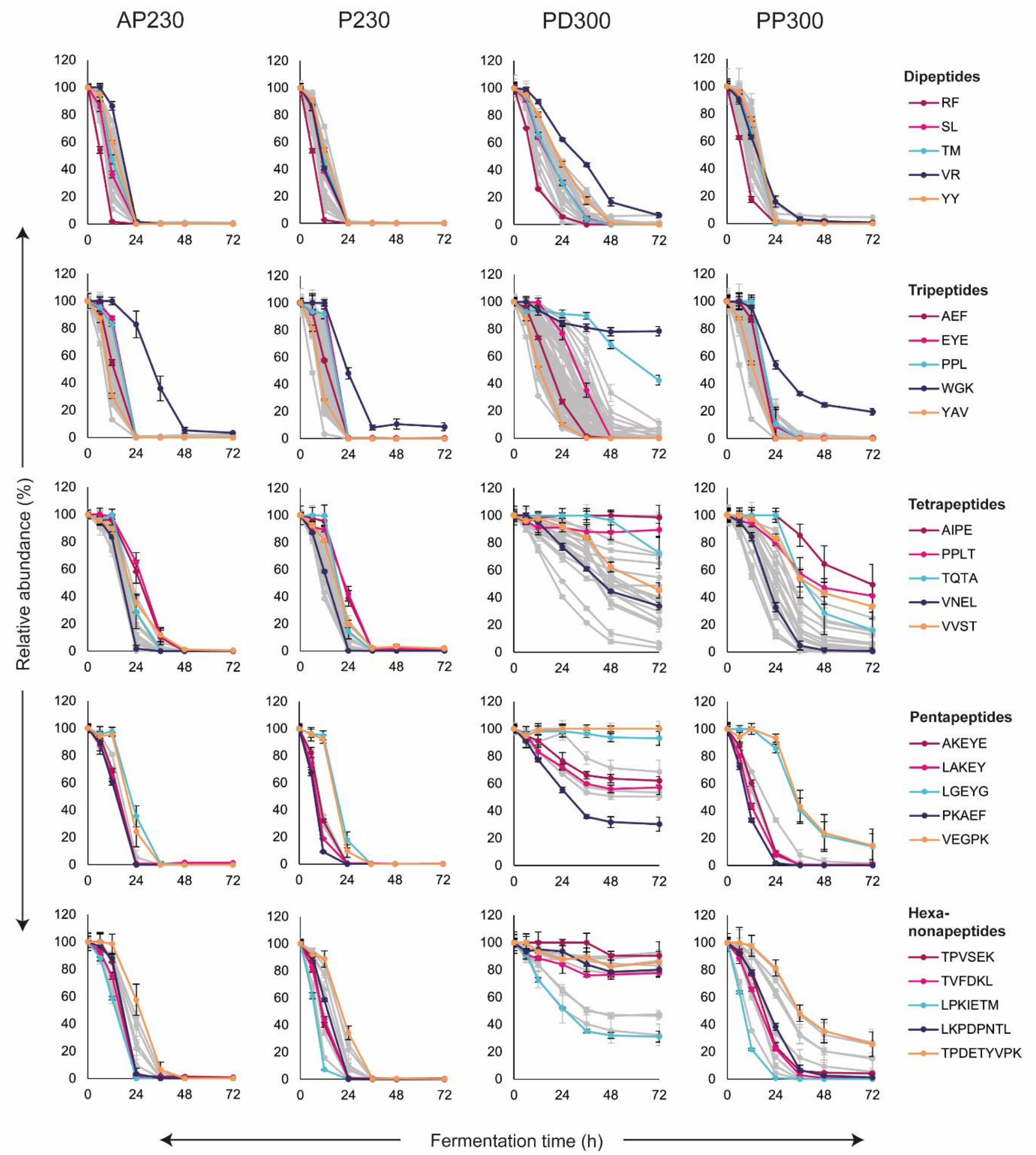


**Figure S1 – Oligopeptide consumption curves of strain SC1 in synthetic whisky mash with different nitrogen conditions.** Five oligopeptides are depicted in colour to represent their peptide length group. All other oligopeptide consumption curves are depicted as 'shadows' in the background.

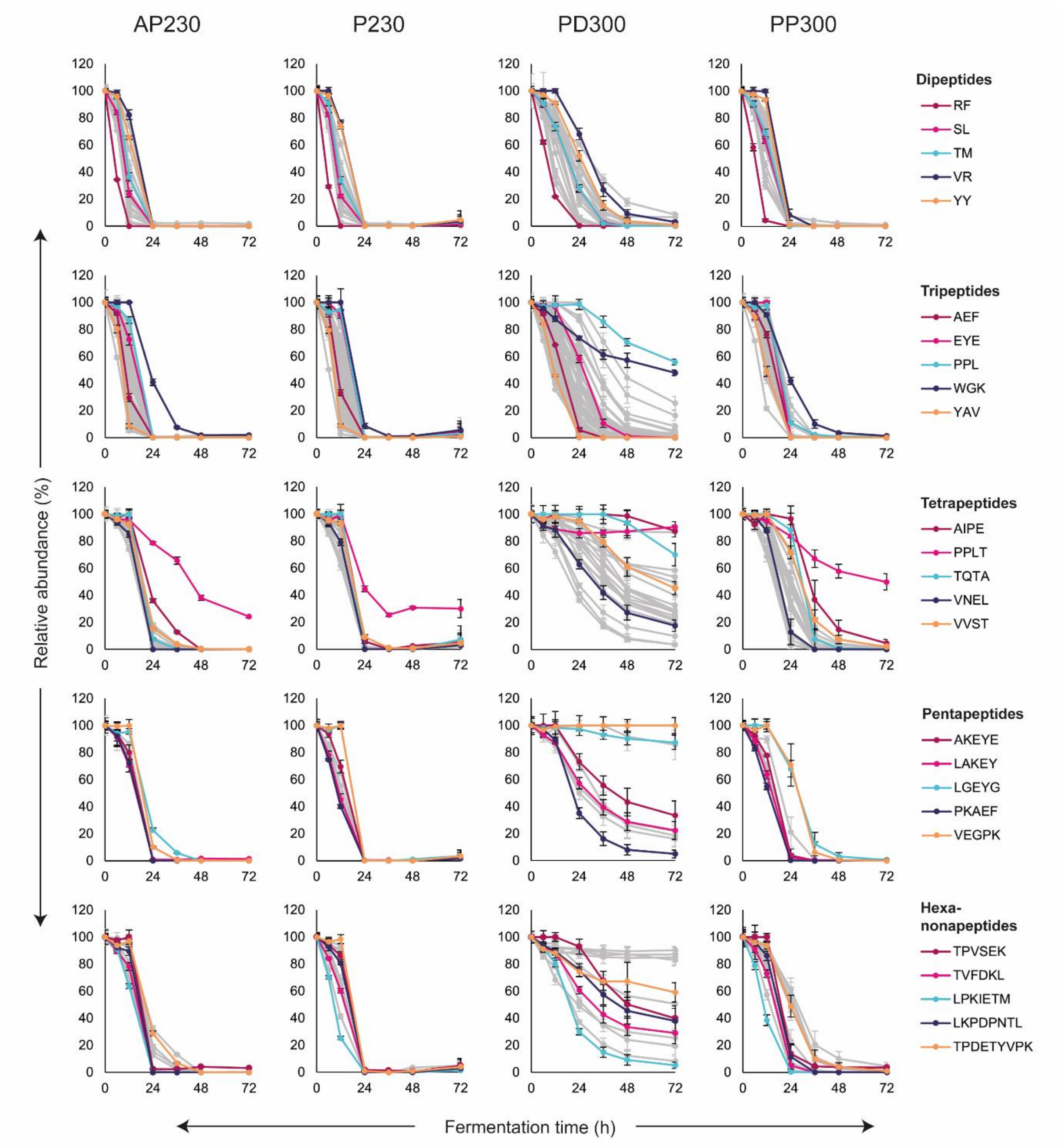


**Figure S2 – Oligopeptide consumption curves of strain SC2 in synthetic whisky mash with different nitrogen conditions.** Five oligopeptides are depicted in colour to represent their peptide length group. All other oligopeptide consumption curves are depicted as 'shadows' in the background.

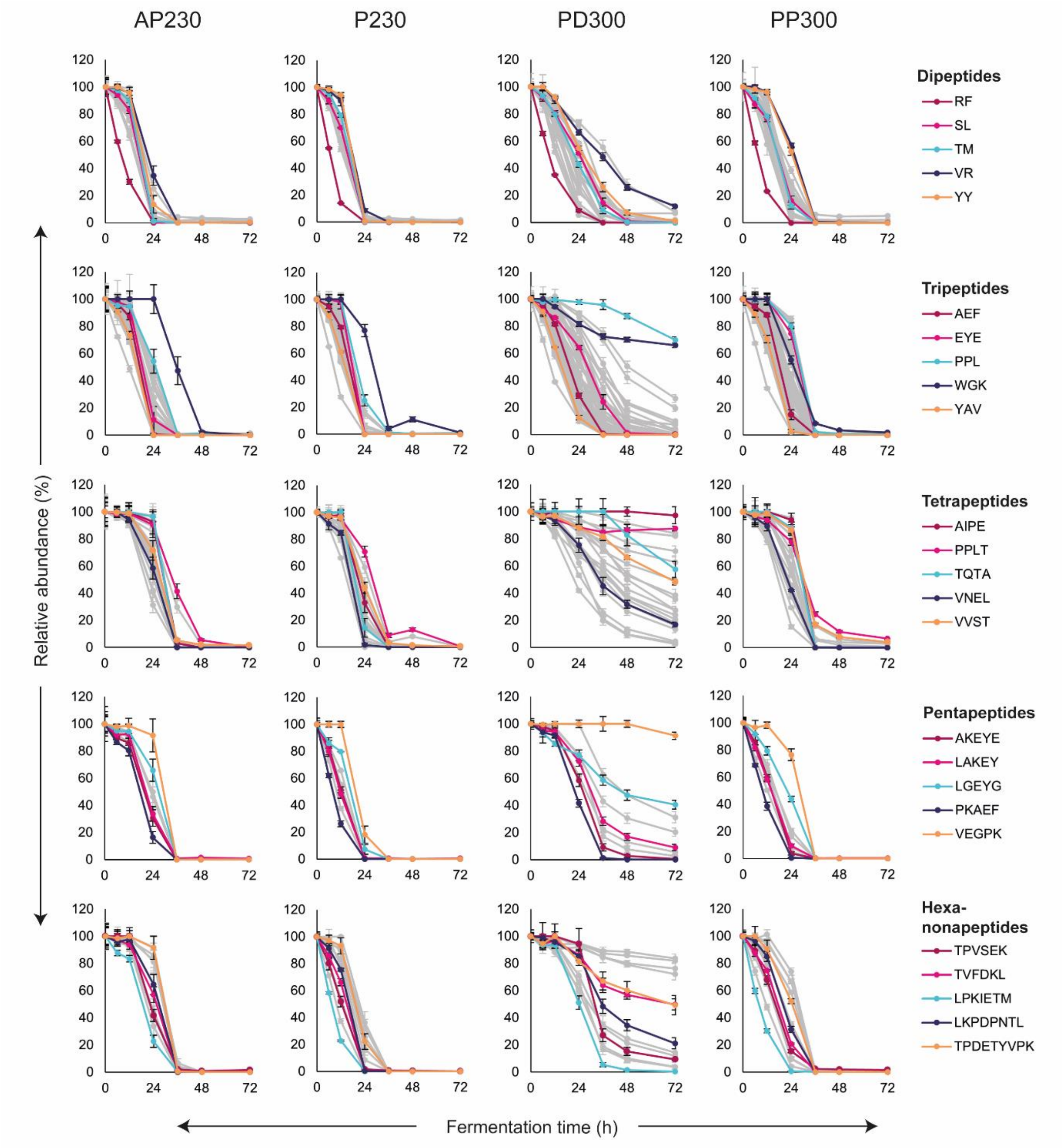


**Figure S3 – Oligopeptide consumption curves of strain SC3 in synthetic whisky mash with different nitrogen conditions.** Five oligopeptides are depicted in colour to represent their peptide length group. All other oligopeptide consumption curves are depicted as ‘shadows’ in the background.

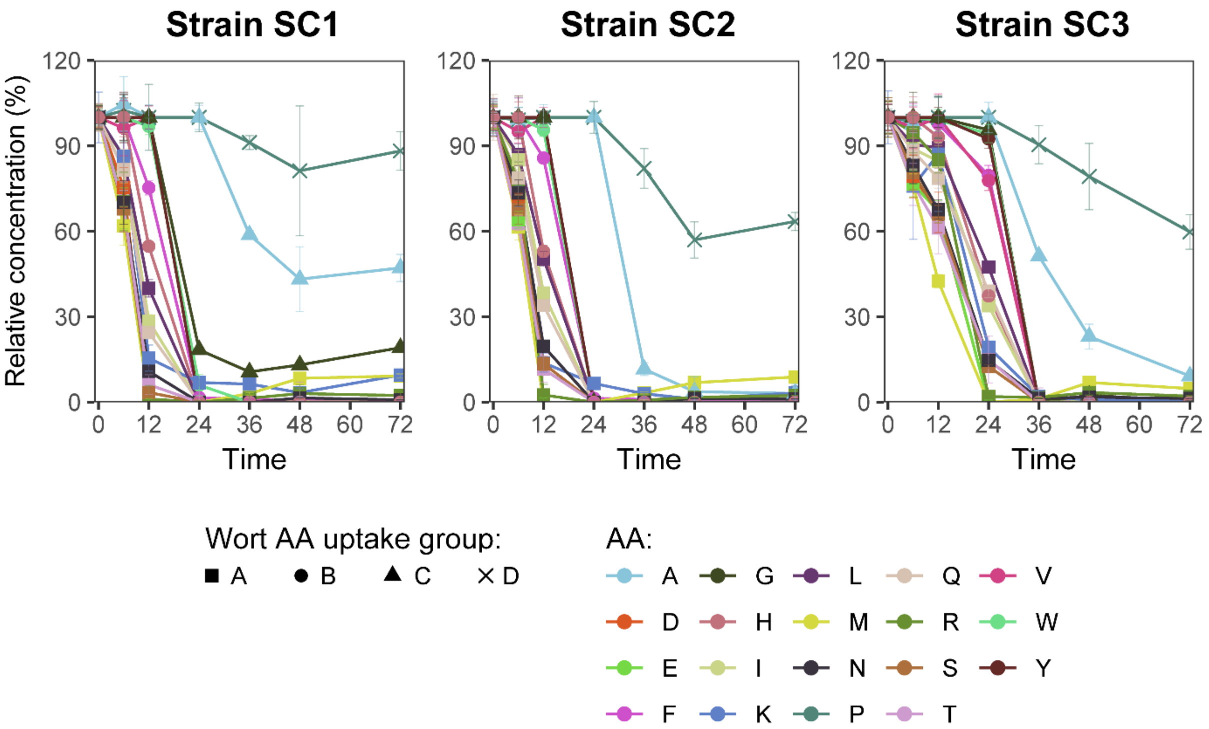


**Figure S4 - Relative amino acid consumption by strains SC1-3 in synthetic whisky mash (AP230).** Individual amino acids have been marked according to their AA uptake order group in wort defined by Jones and Pierce (1964) and revised by Lekkas et al. (2007).

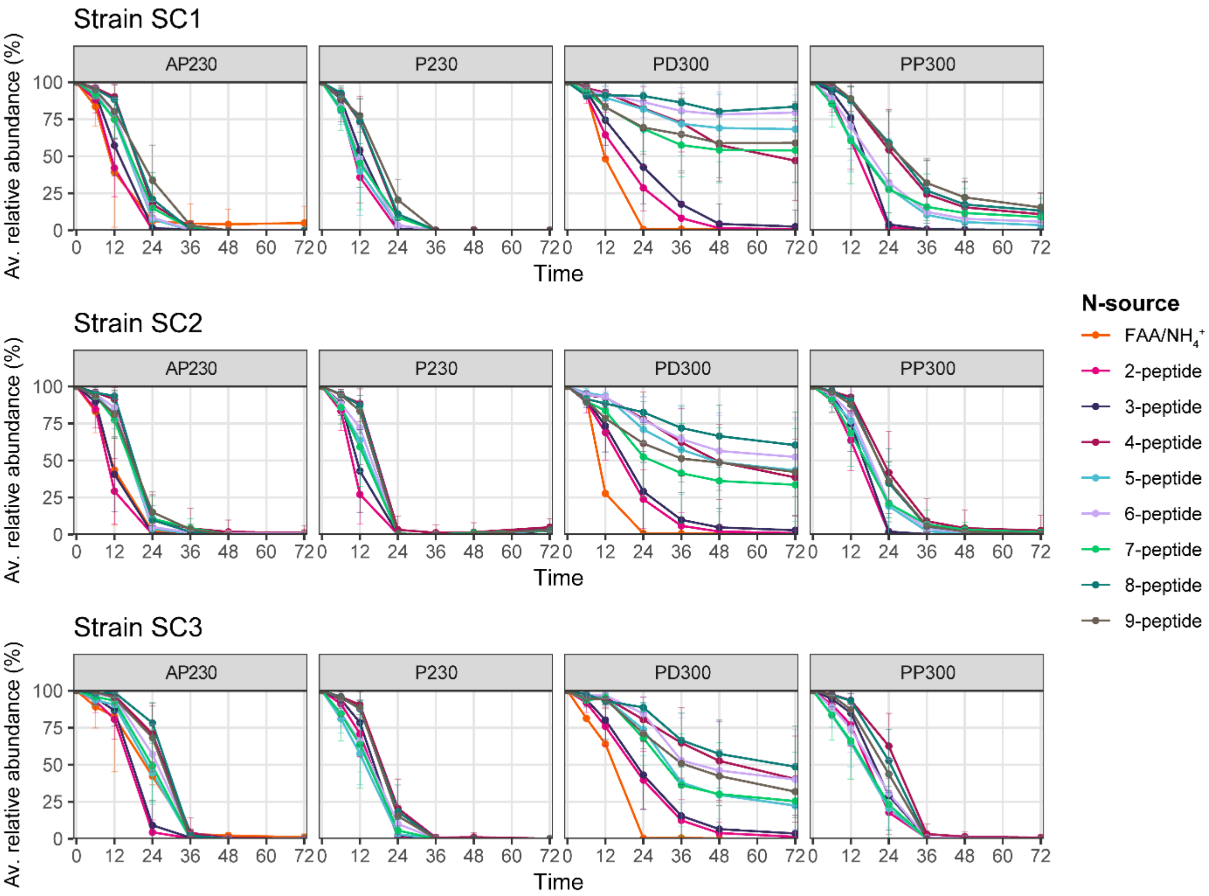


**Figure S5 – Average relative consumption curves sorted by the peptide length group in fermentation with three whisky strains in synthetic whisky mash with different nitrogen conditions.** The orange line in the AP230 medium represents FAA (individual consumption of AAs can be found in Figure S4), while in PD300 it represents ammonium ($NH_4^+$).
